# Identification of a novel and divergent reptarenavirus in an Amazon coral snake (*Micrurus spixii* [Wagler, 1824])

**DOI:** 10.64898/2026.01.29.702531

**Authors:** Ayumu Onishi, Mai Kishimoto, Masayuki Horie

## Abstract

Reptarenaviruses are viruses belonging to the genus *Reptarenavirus* within the family *Arenaviridae*, which infect snakes and cause inclusion body disease (IBD), a fatal condition characterized by behavioral abnormalities and wasting. Although many reptarenaviruses have been identified thus far, the phylogenetic gaps between reptarenaviruses and the other arenaviruses suggest the existence of yet-to-be-identified reptarenaviruses filling the gaps. In this study, we identified a novel reptarenavirus from publicly available RNA-seq data derived from Amazon coral snake (*Micrurus spixii*) and tentatively named it Amazon coral snake virus 1 (ACSV-1). We identified four ACSV-1 contigs containing the putative full-length open reading frames of the NP, GP, and L genes, as well as the partial Z gene. Phylogenetic analyses showed that ACSV-1 is highly divergent from known reptarenaviruses. The NP, GP, and L genes showed 48.3%, 42.3%, and 45.7% nucleotide sequence identities, respectively, with those of the closest relatives. Based on the International Committee on Taxonomy of Viruses (ICTV) species demarcation criteria, ACSV-1 can be assigned to a novel species of virus within the genus *Reptarenavirus*. This study expands our understanding of the diversity and evolution of reptarenaviruses.

## Text

Reptarenaviruses (the genus *Reptarenavirus*) are bi-segmented ambisense RNA viruses within the family *Arenaviridae* [1]. Reptarenaviruses possess two genome segments: the S segment encoding nucleoprotein (NP) and glycoprotein (GP), and the L segment, encoding the large RNA-dependent RNA polymerase (L) and matrix protein (Z). Reptarenaviruses were first identified in 2012 from snakes exhibiting inclusion body disease (IBD), a fatal disease that causes neurological symptoms and wasting, leading to death due to secondary infections [2,3]. After the discoveries, the genus *Reptarenavirus* was established within the family *Arenaviridae* [1]. To date, diverse reptarenaviruses have been detected from captive and wild snakes, and five species are currently assigned in the genus *Reptarenavirus*. However, large phylogenetic gaps remain between reptarenaviruses and other arenaviruses, suggesting that additional reptarenaviruses remain to be discovered. To identify novel arenaviruses, we re-analyzed the data obtained in our previous studies [4,5] and found a contig showing sequence similarity to reptarenavirus L genes in the RNA-seq dataset DRR089660 (data not shown; almost identical to DDBJ YABF01000001). We therefore screened 2,041 RNA-seq datasets derived from the suborder Serpentes (taxid: 8570) in the NCBI SRA database [6] by mapping reads to the above contig using Magic-BLAST [7] (full methods are available in the Supplementary Information). This screening identified six RNA-seq datasets (DRR089660-DRR089665) containing arenavirus-derived reads (Table S1).

To recover viral sequences, raw reads were preprocessed with fastp [8], and host-derived reads were removed by mapping to the genome of *M. fulvius* using HISAT2 [9] and extracting unmapped reads using SAMtools [10]. *De novo* assembly of the unmapped reads was performed using SPAdes [11]. Subsequent BLASTx [12] searches against the NCBI nr database [6] identified contigs with similarity to reptarenavirus NP, GP, L, and Z genes in all six datasets (Tables 1 and S2) (full methods are available in the Supplementary Information). Notably, the number of mapped reads in DRR089663 was substantially higher than in the other five datasets (Table S1). Furthermore, the contig sequences across all six datasets were nearly identical (Table S2), and all datasets originated from the same BioProject (PRJDB5628). These observations strongly suggest that reads in the low-abundance datasets (DRR089660–DRR089662, DRR089664, and DRR089665) likely resulted from index hopping or cross-contamination during library preparation, although low-level infections cannot be completely excluded. Consequently, we conducted further detailed analyses only for the contigs obtained from DRR089663. We validated the accuracy of the contigs obtained from DRR089663 by back-mapping the original reads (Fig. 1A) and trimmed low-confidence regions. The verified contig sequences were deposited in DDBJ under the accession numbers YABF01000001-YABF01000004.

**Table 1.**
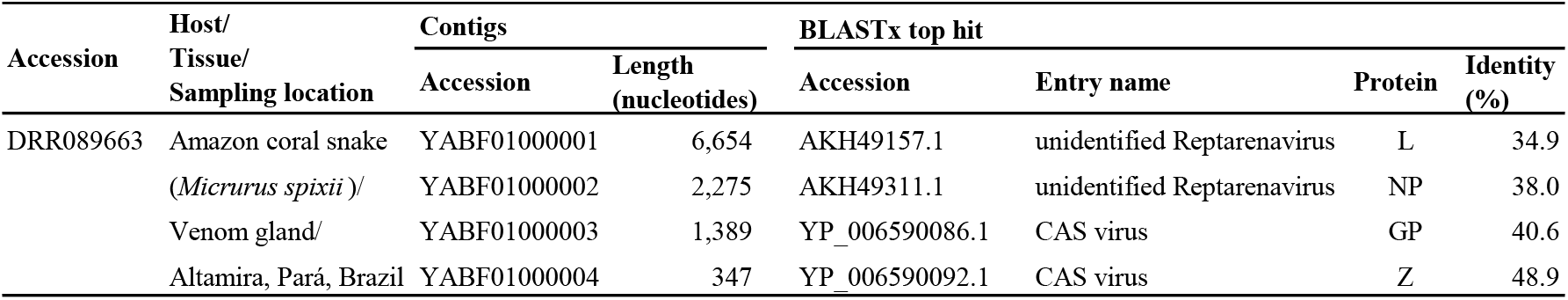
Summary of novel reptarenavirus contigs.

**Figure 1.**
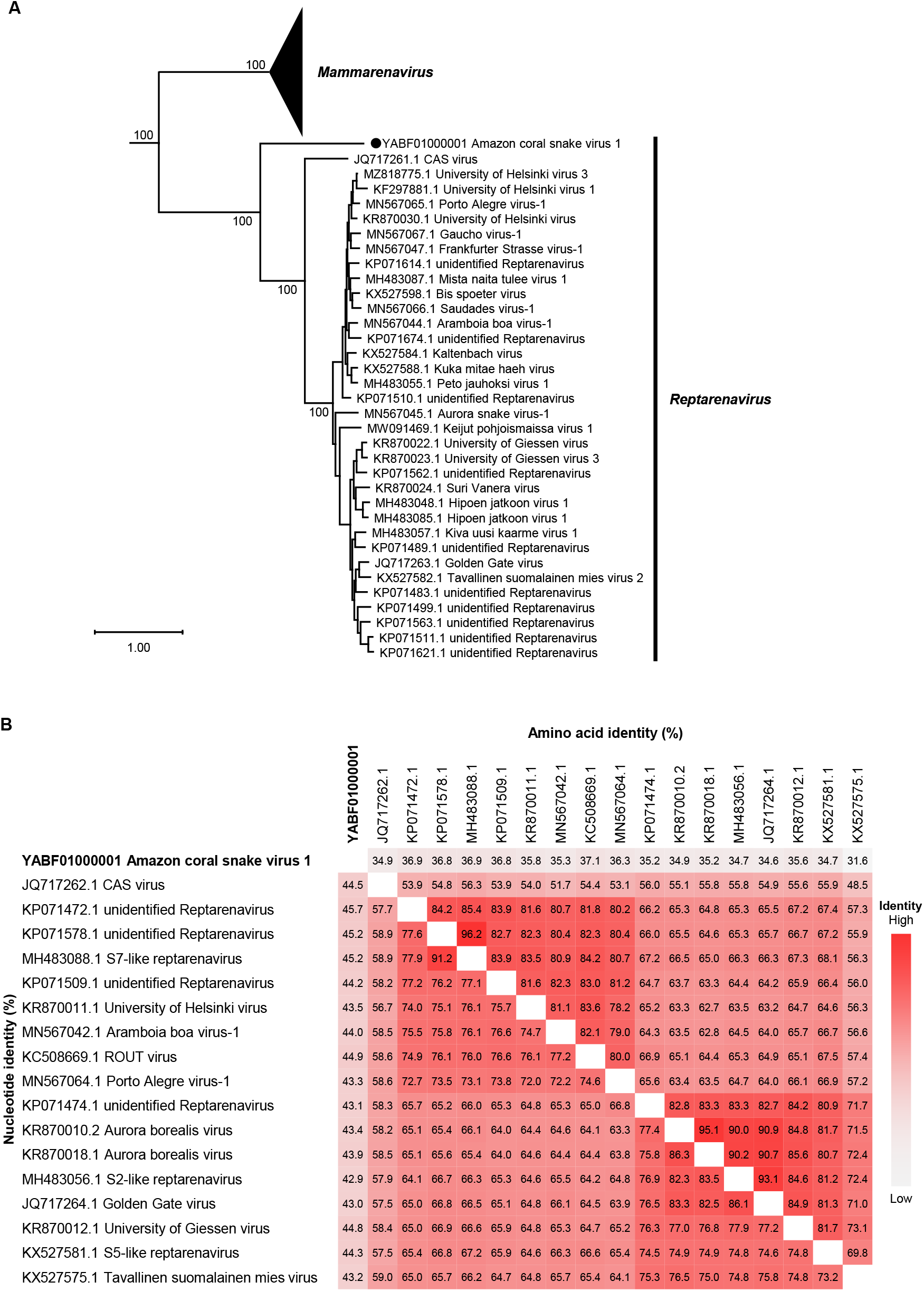
Characterization of Amazon coral snake virus 1. **(A)** Phylogenetic tree of ACSV-1 and representative arenaviruses. The tree was reconstructed by the maximum likelihood method using amino acid sequences of L proteins. Black circle indicates ACSV- 1. Bootstrap values are shown only for key branches. The scale bar indicates the number of amino acid substitutions per site. The full version of the tree is shown in Fig. S3A. **(B)** Pairwise identity heatmap of reptarenavirus NP sequences. Upper and lower triangles indicate nucleotide and amino acid identities, respectively.

We next performed gene annotation of the recovered contigs. The contigs encoding NP, GP, and L appear to contain complete coding sequences, whereas the Z contig does not (Fig. S1; Table 1). Since the reptarenavirus have a bi-segmented genome encoding NP/GP (S segment) and L/Z (L segment), these contigs likely represent transcripts rather than genomic segments. This is most probably due to the use of RACE-seq, which preferentially sequences transcripts [13]. The incomplete recovery of the Z coding sequence may be attributable to the low abundance of Z-derived reads (Table S3). We then performed a series of *in silico* analyses for the putative NP, GP, L, and Z proteins. Domain searches using CD-search [14–16] revealed the putative NP, GP, L, and Z proteins contain Arena_ncap_C, Ebola_HIV-1-like_HR1-HR2, Arena_RNA_pol, and RING_Ubox superfamily domains, respectively, which are conserved among reptarenaviruses (Table S4). SignalP [17] and Deep TMHMM [18] analyses predicted a signal peptide, transmembrane region, and cleavage sites in the GP sequence (Fig. S2).

To gain further insights into the characteristics of ACSV-1, we searched for ACSV-1-derived reads in publicly available RNA-seq datasets from Squamata (taxid: 8509) using all ACSV-1 contigs. We screened 10,208 RNA-seq datasets by mapping, but detected no mappable reads in any datasets other than the six datasets initially identified (Table S3).

To infer the evolutionary relationship between ACSV-1 and other arenaviruses, we performed phylogenetic analyses. First, we reconstructed a phylogenetic tree using the L genes of ACSV-1 and other arenaviruses. The L tree shows that ACSV-1 forms a cluster with members of the genus *Reptarenavirus* but is divergent from known reptarenaviruses (Figs. 1A and S3A). We also reconstructed phylogenetic trees using the NP and GP genes of ACSV-1 and reptarenaviruses. In both trees, ACSV-1 does not form a cluster with any known reptarenaviruses (Figs. S3B and S3C). We further compared nucleotide and amino acid sequences of each ACSV-1 gene with those of representative reptarenaviruses. The nucleotide sequences of the ACSV-1 NP, GP, and L genes showed 48.3%, 42.3%, and 45.7% identities, respectively, with those of the most closely related viruses (Figs. 1B, S4, and S5). The amino acid sequences of the ACSV-1 NP, GP, and L proteins shared 38.3%, 26.8%, and 35.2% identities, respectively, with their closest viral homologs (Figs. 1B, S4, and S5). These results indicate that ACSV-1 is a highly divergent virus within the genus *Reptarenavirus*.

In this study, we identified a genetically divergent reptarenavirus in an Amazon coral snake. To the best of our knowledge, this is the first report of a reptarenavirus detected in a snake belonging to the family *Elapidae*. The current International Committee on Taxonomy of Viruses (ICTV) species demarcation criteria of the genus *Reptarenavirus* are: “(i) the virus shares less than 80% nucleotide sequence identity in the S RNA segment and less than 76% identity in the L RNA segment with other viruses; (ii) association of the virus with a distinct main host or a group of sympatric hosts; (iii) dispersion of the virus in a distinct defined geographical area; and/or (iv) the virus shares less than 88% NP amino-acid sequence identity with other viruses” [1,19]. Here, we showed that ACSV-1 meets criteria (ii) and (iv). Furthermore, although we recovered transcript sequences rather than complete viral genomic segments, the extremely low nucleotide identities between ACSV-1 and known reptarenaviruses strongly suggest that ACSV-1 also meets criterion (i). Thus, we propose that ACSV-1 can be assigned to a novel species within the genus *Reptarenavirus*. Further studies are needed to determine the complete genome sequence.

The pathogenicity of ACSV-1 remains unknown. As described above, some reptarenaviruses have been demonstrated to be causative agents of IBD. Importantly, susceptibility to reptarenaviruses has been reported to differ among host species [20]. Therefore, even if ACSV-1 does not cause obvious disease in Amazon coral snakes, it may pose a risk of IBD-like disease in other snake species. Further epidemiological studies will be necessary to improve our understanding of IBD.

Taken together, this study provides novel insights into the diversity of reptarenaviruses. Further studies are warranted to elucidate the virological characteristics and potential disease association of ACSV-1.

## Author contributions

M.H. conceived and designed the study. All authors performed research and analyzed data. A.O. and M.H. wrote the initial draft. All authors read and approved the final manuscript.

## Acknowledgements

We would like to thank Sakiho Imai for technical advice. The supercomputing resources were provided by the Human Genome Center, the Institute of Medical Science, the University of Tokyo and the NIG supercomputer at ROIS National Institute of Genetics. This study was supported by JSPS KAKENHI grant numbers JP24K21922 (M.H.), JP23K20902 (M.H.), JP22K1923 (M.H.), and JP21H01199 (M.H.) and the Osaka Metropolitan University (OMU) Strategic Research Promotion Project (Young Researcher)(M.K.).

