## Supplementary Figures for "Identification of a novel and divergent reptarenavirus in an Amazon coral snake (*Micrurus spixii* [Wagler, 1824])"

### Supplementary Figure 1

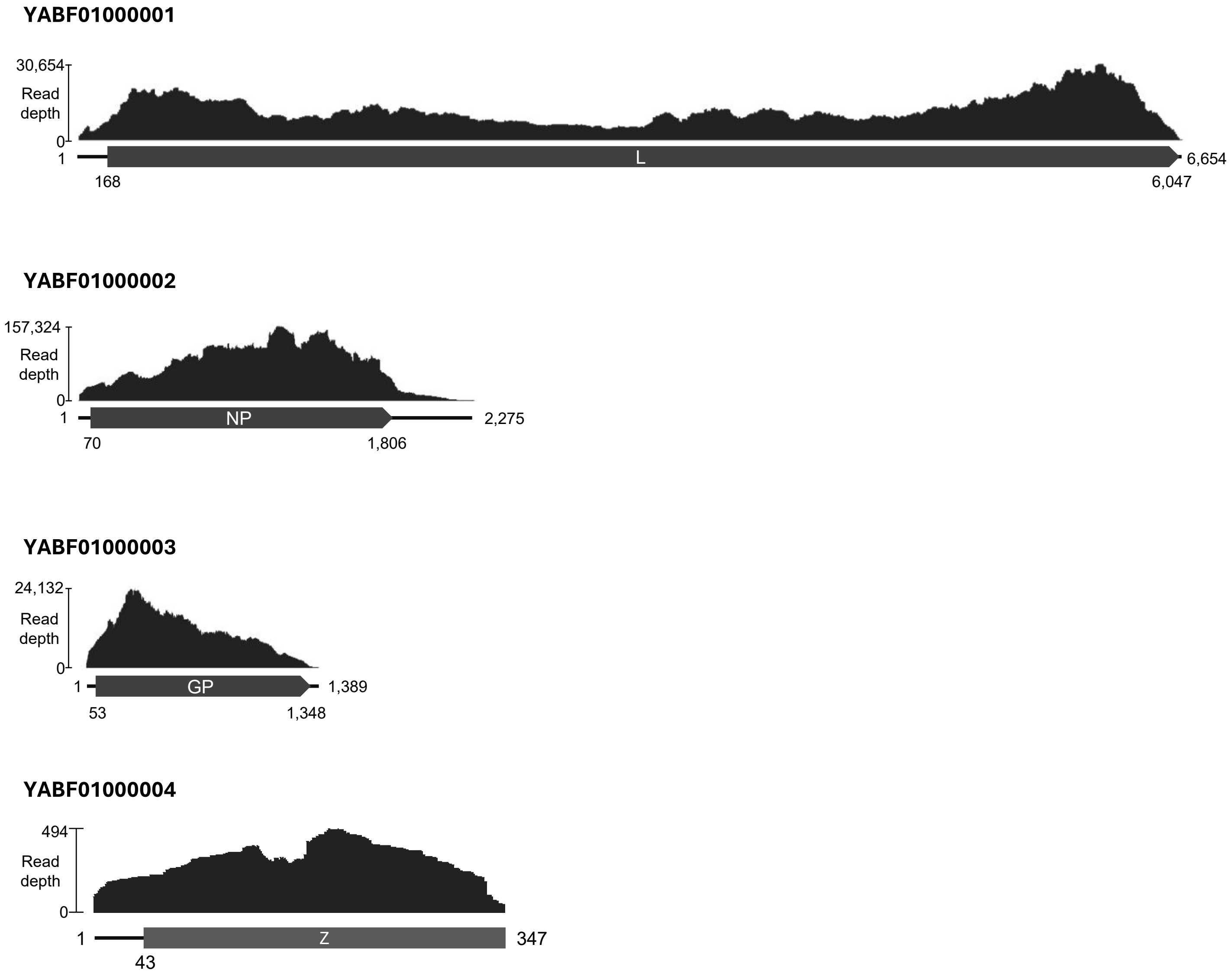


**Supplementary Figure 1. Identified ACSV-1 contigs and read-depth profiles**. Short reads from DRR089663 were mapped to the ACSV-1 contigs, and per-base read depth was visualized. Gray boxes indicate open reading frames (ORFs). Numbers represent nucleotide positions on the contigs. Note that the y-axis scale for the Z contig differs from those of the other contigs.

### Supplementary Figure 2

**A**

Cleavage site between pos. 22 and 23.
Probability 0.965263

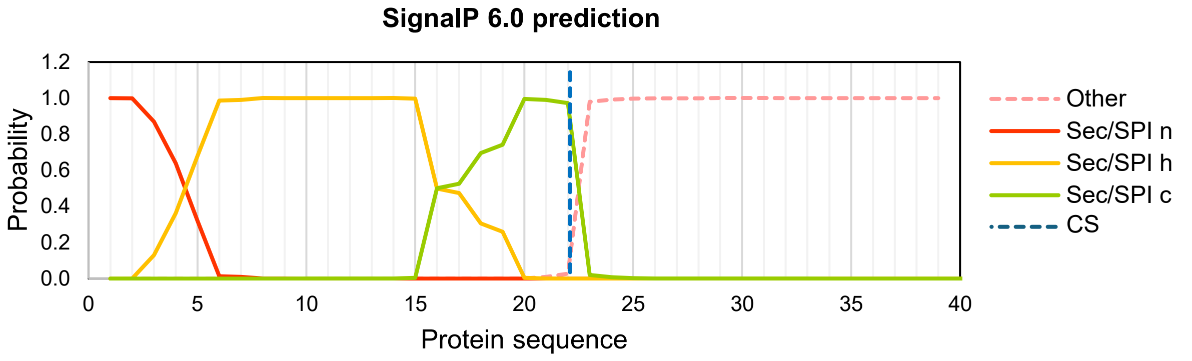


**B**

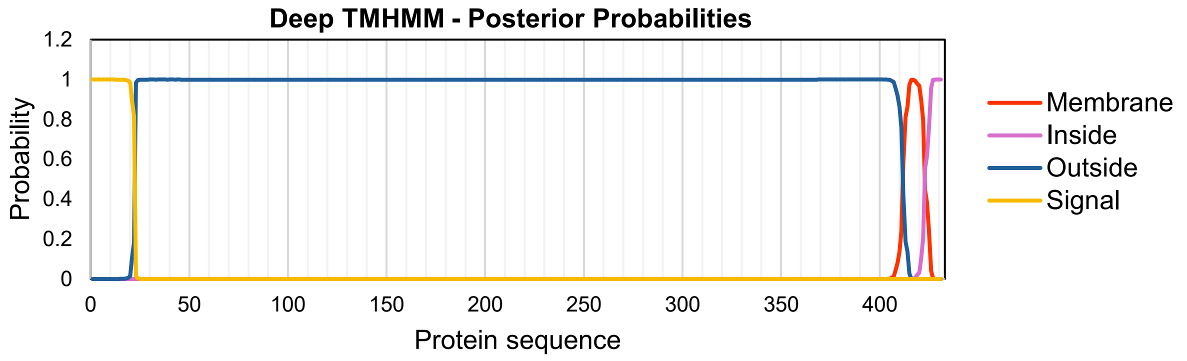


**Supplementary Figure 2. *In silico* characterization of GP of Amazon coral snake virus 1.**
**(A)** Signal peptide prediction was performed using the SignalP 6.0 server. Labels n, h, and c represent the predicted N-terminal region, central hydrophobic region, and C-terminal region of signal peptide, respectively. CS indicates the predicted cleavage site. **(B)** Transmembrane region was predicted using the DeepTMHMM 1.0 server. Predicted regions: signal peptide (1–22), extracellular domain (23–411), transmembrane domain (412–422), and cytoplasmic tail (423–431).

### Supplementary Figure 3

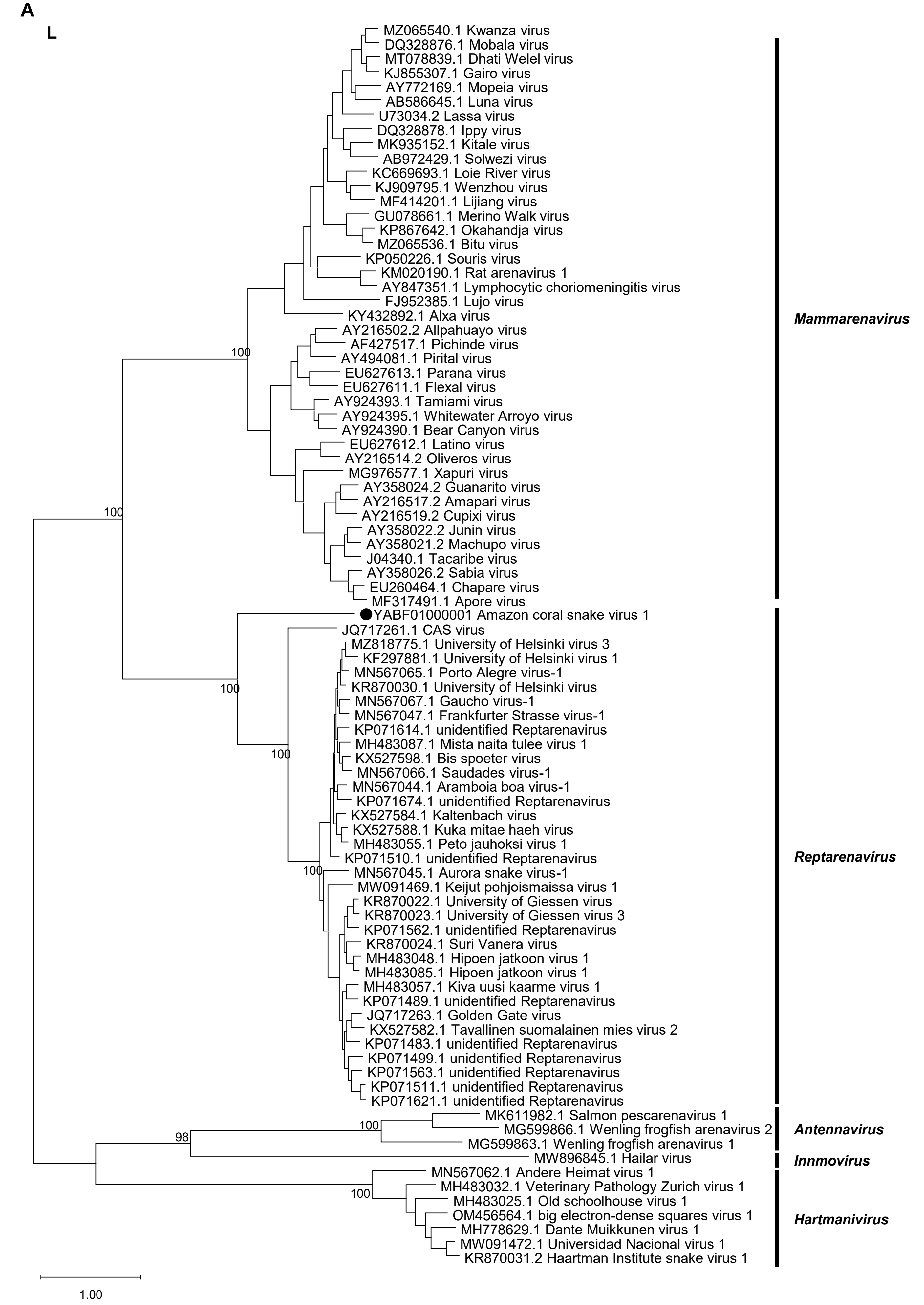

### Supplementary Figure 3 (continued)

B

NP

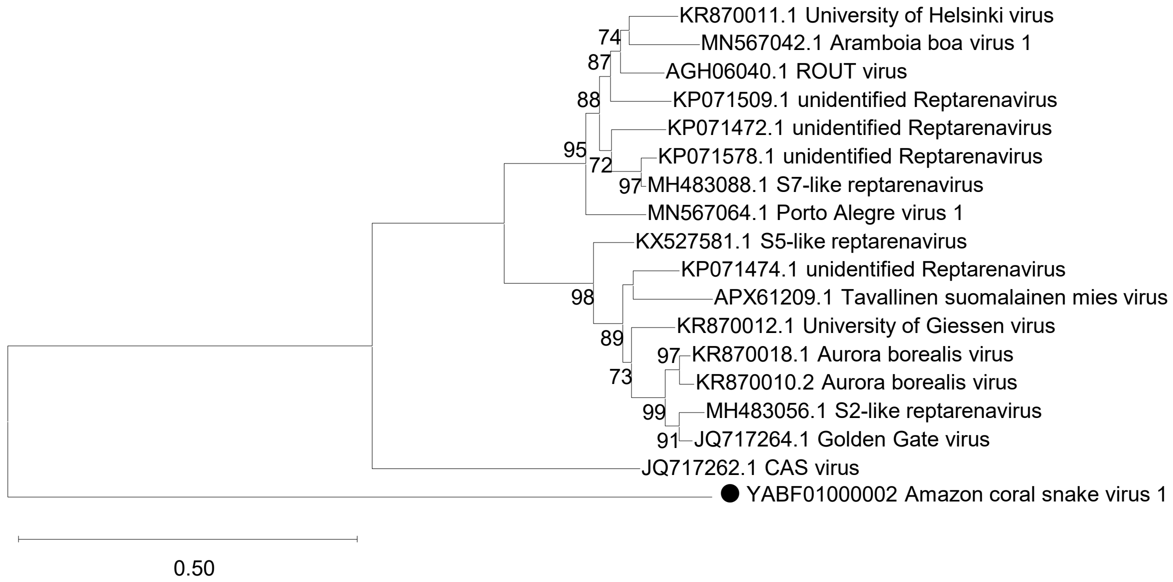


C

GP

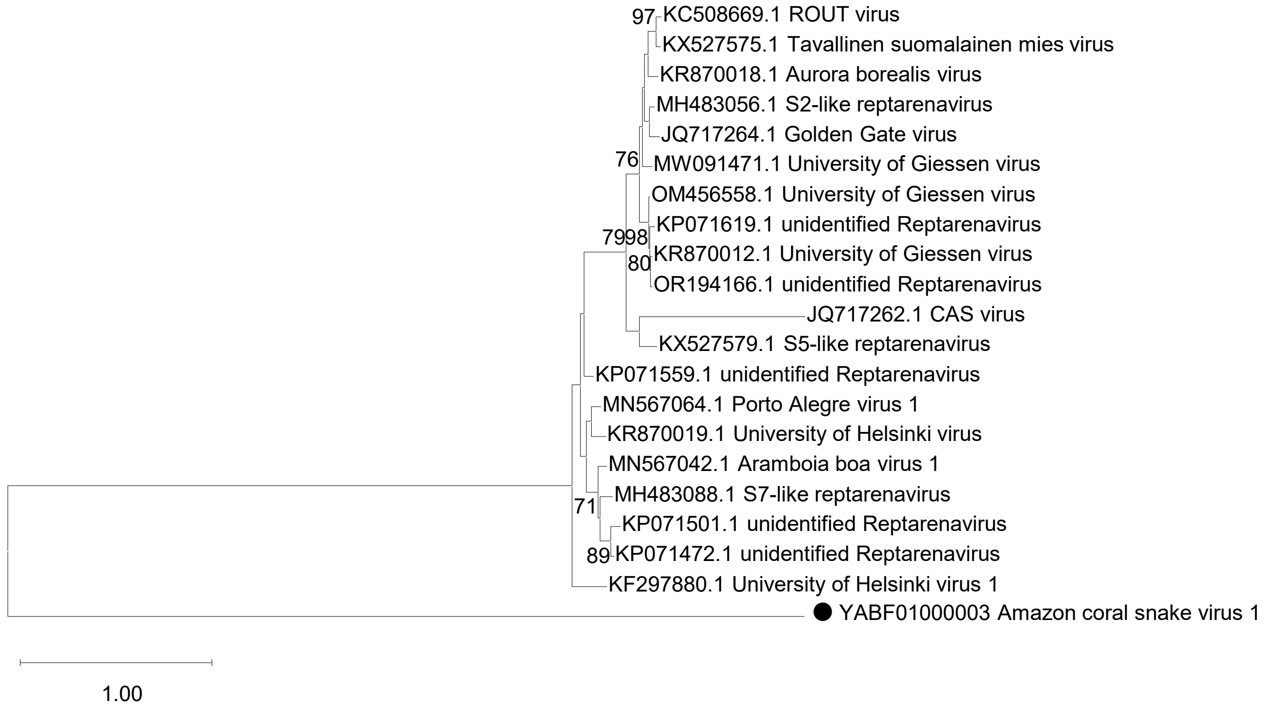


**Supplementary Figure 3. Phylogenetic relationship of Amazon coral snake virus 1 and other arenaviruses.** Phylogenetic tree of Amazon coral snake virus 1 (ACSV-1) and arenaviruses reconstructed by the maximum likelihood method using amino acid sequences of L **(A)**, NP **(B)**, and GP **(C)** proteins. ACSV-1 is indicated with a black circle. Bootstrap values more than 70 are shown on each branch. The scale bar represents the number of amino acid substitutions per site.

Amino acid identity (%)

Nucleotide identity (%)

| | **YABF01000003** | JQ717262.1 | KP071472.1 | KP071501.1 | MN567042.1 | MH483088.1 | KP071559.1 | KF297880.1 | KR870019.1 | MN567064.1 | KP071619.1 | KR870012.1 | OR194166.1 | OM456558.1 | KR870018.1 | KC508669.1 | KX527575.1 | MH483056.1 | JQ717264.1 | MW091471.1 | KX527579.1 |
|---|---|---|---|---|---|---|---|---|---|---|---|---|---|---|---|---|---|---|---|---|---|
| **YABF01000003 Amazon coral snake virus 1** | | 21.5 | 22.3 | 22.3 | 23.4 | 24.2 | 23.5 | 23.7 | 23.9 | 23.7 | 22.4 | 22.3 | 22.3 | 22.7 | 22.4 | 21.9 | 22.6 | 23.6 | 22.6 | 21.9 | 23.0 |
| JQ717262.1 CAS virus | 33.0 | | 47.5 | 47.5 | 47.0 | 45.7 | 47.4 | 43.4 | 47.7 | 45.9 | 46.6 | 46.0 | 46.2 | 47.3 | 47.1 | 46.3 | 46.9 | 47.9 | 47.2 | 47.8 | 46.9 |
| KP071472.1 unidentified Reptarenavirus | 36.8 | 48.7 | | 93.7 | 88.4 | 85.3 | 74.5 | 69.0 | 78.7 | 78.0 | 64.8 | 64.3 | 63.6 | 64.5 | 64.5 | 63.6 | 63.8 | 65.5 | 65.0 | 64.0 | 62.1 |
| KP071501.1 unidentified Reptarenavirus | 36.2 | 48.3 | 94.2 | | 86.7 | 83.6 | 74.2 | 69.7 | 76.8 | 77.0 | 66.7 | 65.9 | 65.7 | 66.8 | 65.7 | 65.5 | 66.1 | 66.4 | 65.5 | 63.8 | 63.3 |
| MN567042.1 Aramboia boa virus-1 | 35.9 | 48.8 | 80.1 | 78.6 | | 85.3 | 76.4 | 69.5 | 78.5 | 79.4 | 64.8 | 64.5 | 64.3 | 65.7 | 65.4 | 64.8 | 65.0 | 66.2 | 66.0 | 65.9 | 63.6 |
| MH483088.1 S7-like reptarenavirus | 36.7 | 49.2 | 78.2 | 76.5 | 77.8 | | 74.2 | 67.8 | 76.8 | 78.0 | 65.5 | 64.5 | 64.5 | 65.7 | 65.4 | 65.0 | 65.2 | 65.5 | 64.6 | 65.4 | 63.3 |
| KP071559.1 unidentified Reptarenavirus | 34.6 | 48.7 | 68.2 | 67.6 | 69.2 | 68.6 | | 76.1 | 77.5 | 79.0 | 65.2 | 64.2 | 64.4 | 64.9 | 65.8 | 64.7 | 65.3 | 66.8 | 65.7 | 65.1 | 65.3 |
| KF297880.1 University of Helsinki virus 1 | 33.6 | 47.7 | 67.6 | 67.2 | 68.3 | 66.9 | 75.6 | | 71.2 | 72.8 | 60.3 | 60.0 | 59.4 | 60.5 | 61.2 | 59.4 | 60.5 | 61.7 | 61.7 | 59.8 | 59.1 |
| KR870019.1 University of Helsinki virus | 35.4 | 47.8 | 70.5 | 69.8 | 71.8 | 70.9 | 71.8 | 70.9 | | 84.4 | 63.1 | 63.0 | 62.8 | 63.5 | 64.9 | 63.3 | 63.7 | 64.3 | 65.0 | 64.2 | 62.6 |
| MN567064.1 Porto Alegre virus-1 | 35.1 | 48.8 | 71.0 | 70.9 | 72.1 | 71.6 | 73.4 | 72.8 | 76.6 | | 65.0 | 64.2 | 64.2 | 65.1 | 66.7 | 64.7 | 65.3 | 65.2 | 64.5 | 65.4 | 64.7 |
| KP071619.1 unidentified Reptarenavirus | 37.1 | 50.3 | 61.8 | 64.9 | 62.9 | 62.9 | 62.2 | 61.4 | 61.3 | 62.2 | | 95.1 | 93.7 | 94.9 | 83.6 | 82.8 | 82.7 | 83.2 | 83.0 | 84.8 | 74.8 |
| KR870012.1 University of Giessen virus 1 | 36.9 | 50.1 | 61.4 | 64.7 | 62.6 | 62.2 | 61.6 | 60.8 | 61.8 | 61.7 | 96.7 | | 95.6 | 94.6 | 82.1 | 82.3 | 82.1 | 83.4 | 83.7 | 84.6 | 74.4 |
| OR194166.1 Reptarenavirus sp. | 37.2 | 50.3 | 61.4 | 64.5 | 62.2 | 62.1 | 61.4 | 60.6 | 61.6 | 61.4 | 95.8 | 97.4 | | 94.4 | 82.8 | 82.5 | 81.8 | 83.2 | 83.7 | 85.5 | 75.3 |
| OM456558.1 University of Giessen virus | 37.3 | 50.0 | 61.9 | 64.8 | 63.1 | 62.2 | 62.4 | 61.7 | 61.3 | 61.8 | 94.3 | 94.3 | 94.3 | | 83.2 | 83.7 | 83.9 | 84.6 | 86.0 | 86.7 | 74.8 |
| KR870018.1 Aurora borealis virus | 37.7 | 49.7 | 62.1 | 63.8 | 62.3 | 63.0 | 61.9 | 61.9 | 59.8 | 63.7 | 75.3 | 75.2 | 75.3 | 75.4 | | 85.9 | 84.5 | 84.1 | 83.4 | 84.6 | 73.8 |
| KC508669.1 ROUT virus | 35.3 | 49.3 | 60.0 | 61.9 | 61.4 | 61.3 | 62.2 | 62.2 | 60.7 | 62.6 | 75.9 | 75.1 | 75.0 | 76.7 | 81.5 | | 92.5 | 84.1 | 82.4 | 83.4 | 73.1 |
| KX527575.1 Tavallinen suomalainen mies virus 2 | 36.2 | 48.8 | 60.7 | 62.5 | 61.9 | 61.1 | 62.6 | 62.2 | 60.4 | 62.0 | 75.6 | 74.9 | 74.8 | 76.7 | 81.1 | 90.5 | | 82.9 | 82.2 | 81.8 | 74.3 |
| MH483056.1 S2-like reptarenavirus | 37.5 | 49.9 | 61.8 | 63.0 | 62.0 | 61.7 | 62.2 | 61.5 | 60.1 | 62.2 | 75.2 | 75.0 | 74.9 | 76.0 | 79.2 | 79.0 | 79.7 | | 88.5 | 86.7 | 76.2 |
| JQ717264.1 Golden Gate virus | 36.6 | 49.3 | 61.1 | 62.5 | 62.0 | 62.7 | 61.6 | 62.9 | 60.2 | 62.4 | 75.7 | 75.5 | 76.3 | 77.8 | 77.6 | 78.9 | 78.9 | 83.9 | | 86.7 | 74.5 |
| MW091471.1 University of Giessen virus | 36.2 | 50.4 | 62.7 | 63.4 | 62.3 | 62.5 | 61.4 | 61.2 | 60.7 | 61.8 | 78.3 | 77.9 | 77.9 | 79.1 | 78.1 | 78.3 | 77.8 | 78.6 | 79.9 | | 74.8 |
| KX527579.1 S5-like reptarenavirus | 37.5 | 48.7 | 61.4 | 61.6 | 63.0 | 61.7 | 63.4 | 61.7 | 61.0 | 64.0 | 70.4 | 70.2 | 70.5 | 70.2 | 68.7 | 68.5 | 69.1 | 70.1 | 69.3 | 69.5 | |

**Supplementary Figure 4. Pairwise identity heatmap of reptarenavirus GP sequences.** Pairwise identity heatmap of reptarenavirus GP sequences are shown. Upper and lower triangles indicate nucleotide and amino acid identities, respectively.

Amino acid identity (%)

Nucleotide identity (%)

| | **YABF01000001** | JQ717261.1 | KP071483.1 | KP071489.1 | KR870028.1 | JQ717263.1 | KX527582.1 | KP071499.1 | KP071563.1 | KP071621.1 | KP071562.1 | MH483051.1 | MH483048.1 | KR870024.1 | KP071510.1 | KP071548.1 | KR870020.1 | KF297881.1 | MN567067.1 | KX527598.1 | MN567044.1 | KX527588.1 | KP071674.1 | MW091469.1 | MN567045.1 |
|---|---|---|---|---|---|---|---|---|---|---|---|---|---|---|---|---|---|---|---|---|---|---|---|---|---|
| **YABF01000001 Amazon coral snake virus 1** | | 32.7 | 33.2 | 32.5 | 31.9 | 32.3 | 31.8 | 33.1 | 33.0 | 33.2 | 31.9 | 31.6 | 32.6 | 32.6 | 32.2 | 31.7 | 32.2 | 32.6 | 31.5 | 31.9 | 32.2 | 31.7 | 31.8 | 31.7 | 32.3 |
| JQ717261.1 CAS virus | 40.3 | | 48.8 | 48.9 | 48.7 | 49.0 | 48.0 | 47.8 | 49.0 | 48.7 | 47.6 | 48.0 | 47.0 | 48.2 | 47.6 | 48.2 | 48.3 | 47.7 | 48.0 | 48.3 | 48.3 | 47.9 | 47.8 | 48.6 | 49.5 |
| KP071483.1 unidentified Reptarenavirus | 43.0 | 54.4 | | 76.8 | 76.8 | 76.3 | 74.3 | 75.5 | 74.8 | 74.1 | 74.0 | 74.3 | 72.4 | 73.1 | 64.6 | 64.4 | 65.0 | 67.7 | 64.4 | 65.1 | 65.1 | 64.3 | 64.9 | 65.1 | 65.4 |
| KP071489.1 unidentified Reptarenavirus | 42.5 | 53.7 | 72.4 | | 77.1 | 75.7 | 74.1 | 75.4 | 74.8 | 73.8 | 74.0 | 74.5 | 72.1 | 72.8 | 64.9 | 64.1 | 64.2 | 66.6 | 64.5 | 64.5 | 64.7 | 63.9 | 64.3 | 65.3 | 65.1 |
| KR870028.1 Hans Kompis virus | 42.6 | 54.2 | 72.4 | 71.5 | | 76.0 | 74.7 | 74.0 | 74.2 | 73.2 | 75.0 | 73.9 | 72.8 | 72.9 | 65.1 | 64.2 | 64.6 | 67.5 | 64.4 | 64.1 | 64.9 | 64.5 | 64.6 | 64.8 | 65.2 |
| JQ717263.1 Golden Gate virus | 42.7 | 54.4 | 71.7 | 71.0 | 71.7 | | 77.5 | 74.9 | 74.6 | 74.2 | 73.1 | 72.8 | 72.8 | 72.9 | 64.7 | 64.4 | 64.1 | 71.2 | 64.1 | 63.7 | 64.4 | 64.3 | 64.8 | 66.2 | 65.6 |
| KX527582.1 Tavallinen suomalainen mies virus 2 | 42.2 | 54.3 | 70.4 | 70.0 | 70.9 | 72.0 | | 72.7 | 73.4 | 73.0 | 72.1 | 72.6 | 71.4 | 72.4 | 62.8 | 63.5 | 63.3 | 66.8 | 63.3 | 62.5 | 63.3 | 63.3 | 63.4 | 64.1 | 63.5 |
| KP071499.1 unidentified Reptarenavirus | 42.7 | 53.9 | 71.0 | 70.9 | 70.4 | 70.0 | 68.7 | | 76.3 | 75.3 | 72.1 | 71.9 | 71.4 | 71.0 | 62.8 | 63.2 | 63.3 | 66.6 | 62.3 | 62.7 | 63.0 | 63.2 | 63.0 | 63.4 | 63.4 |
| KP071563.1 unidentified Reptarenavirus | 42.6 | 54.5 | 70.8 | 70.9 | 70.8 | 71.3 | 69.7 | 72.0 | | 78.1 | 72.6 | 72.0 | 71.6 | 71.7 | 63.5 | 64.0 | 63.9 | 66.2 | 63.8 | 63.6 | 63.9 | 63.3 | 63.7 | 64.9 | 63.9 |
| KP071621.1 unidentified Reptarenavirus | 42.6 | 54.4 | 70.7 | 70.7 | 70.0 | 71.3 | 69.2 | 72.1 | 73.5 | | 70.6 | 71.0 | 70.0 | 70.9 | 63.2 | 63.7 | 63.7 | 66.0 | 63.0 | 62.7 | 64.2 | 63.9 | 63.2 | 64.5 | 62.7 |
| KP071562.1 unidentified Reptarenavirus | 41.9 | 53.8 | 70.3 | 70.4 | 70.9 | 69.7 | 68.9 | 68.6 | 69.3 | 68.2 | | 79.5 | 76.7 | 75.5 | 64.5 | 63.7 | 64.3 | 65.7 | 64.0 | 64.0 | 64.0 | 63.7 | 64.2 | 64.7 | 64.6 |
| MH483051.1 University of Giessen virus 1 | 42.5 | 53.8 | 70.2 | 70.1 | 70.6 | 69.7 | 68.7 | 68.7 | 68.6 | 68.1 | 74.5 | | 76.9 | 76.3 | 64.0 | 63.5 | 64.1 | 65.3 | 63.2 | 64.0 | 64.3 | 63.9 | 64.5 | 65.2 | 63.6 |
| MH483048.1 Hipoen jatkoon virus 1 | 42.0 | 53.1 | 69.1 | 68.5 | 69.0 | 68.9 | 67.9 | 68.1 | 68.2 | 67.3 | 72.2 | 72.7 | | 76.1 | 63.6 | 64.5 | 64.2 | 65.7 | 63.3 | 64.1 | 64.7 | 63.7 | 63.9 | 64.5 | 64.4 |
| KR870024.1 Suri Vanera virus | 42.5 | 53.6 | 69.7 | 69.4 | 69.5 | 68.6 | 68.9 | 68.4 | 68.1 | 68.1 | 71.8 | 72.2 | 71.4 | | 65.1 | 64.2 | 64.6 | 65.6 | 64.0 | 64.5 | 63.9 | 64.2 | 64.6 | 65.3 | 63.7 |
| KP071510.1 unidentified Reptarenavirus | 41.7 | 53.2 | 63.2 | 62.7 | 63.7 | 63.1 | 62.5 | 61.7 | 63.1 | 61.9 | 62.9 | 62.4 | 61.9 | 62.6 | | 78.5 | 80.6 | 73.4 | 78.3 | 78.7 | 78.4 | 78.1 | 75.8 | 64.8 | 67.4 |
| KP071548.1 unidentified Reptarenavirus | 41.3 | 53.0 | 63.1 | 62.4 | 62.7 | 62.9 | 62.2 | 62.7 | 62.6 | 62.3 | 62.5 | 62.4 | 62.4 | 62.4 | 72.1 | | 80.0 | 72.8 | 78.1 | 78.3 | 78.4 | 77.6 | 76.2 | 64.3 | 65.6 |
| KR870020.1 University of Helsinki virus | 42.3 | 53.4 | 63.7 | 63.0 | 63.4 | 62.9 | 62.3 | 62.4 | 62.2 | 62.9 | 63.4 | 63.6 | 62.3 | 62.6 | 73.5 | 73.5 | | 87.5 | 81.4 | 81.5 | 80.6 | 80.3 | 77.2 | 65.9 | 66.4 |
| KF297881.1 University of Helsinki virus 1 | 41.9 | 53.4 | 65.8 | 65.1 | 66.2 | 68.2 | 65.0 | 65.0 | 64.3 | 65.3 | 65.1 | 65.1 | 64.0 | 64.0 | 69.1 | 69.6 | 89.7 | | 73.6 | 74.4 | 73.0 | 73.2 | 71.7 | 64.3 | 64.3 |
| MN567067.1 Gaucho virus-1 | 42.1 | 53.0 | 62.9 | 63.0 | 62.9 | 62.0 | 62.1 | 62.1 | 62.7 | 62.3 | 63.3 | 62.3 | 61.8 | 62.2 | 72.5 | 73.2 | 74.6 | 70.1 | | 78.8 | 77.5 | 78.6 | 75.8 | 65.1 | 65.9 |
| KX527598.1 Bis spoeter virus | 41.1 | 53.0 | 62.8 | 61.8 | 62.7 | 62.1 | 61.0 | 61.2 | 62.6 | 60.9 | 62.0 | 62.0 | 61.5 | 61.8 | 73.0 | 73.2 | 73.2 | 69.1 | 72.8 | | 77.7 | 78.7 | 76.0 | 64.0 | 65.5 |
| MN567044.1 Aramboia boa virus-1 | 42.2 | 53.3 | 63.6 | 63.0 | 64.0 | 63.1 | 62.2 | 62.0 | 62.8 | 62.8 | 63.2 | 62.9 | 62.7 | 62.2 | 72.9 | 72.9 | 73.6 | 69.9 | 72.6 | 72.7 | | 77.6 | 76.6 | 64.7 | 65.5 |
| KX527588.1 Kuka mitae haeh virus | 41.4 | 52.6 | 62.8 | 62.6 | 62.7 | 62.8 | 61.5 | 62.2 | 62.6 | 62.2 | 62.4 | 62.6 | 62.0 | 62.1 | 72.1 | 73.3 | 72.7 | 68.5 | 72.3 | 73.4 | 72.3 | | 75.7 | 65.1 | 65.2 |
| KP071674.1 unidentified Reptarenavirus | 42.1 | 52.3 | 63.3 | 61.9 | 62.9 | 63.1 | 62.0 | 61.5 | 62.0 | 61.8 | 62.2 | 62.6 | 62.3 | 62.6 | 70.7 | 70.9 | 71.4 | 68.2 | 70.7 | 70.7 | 71.8 | 69.7 | | 65.2 | 65.2 |
| MW091469.1 Keijut pohjoismaissa virus 1 | 41.8 | 52.8 | 63.4 | 64.3 | 64.1 | 63.7 | 63.1 | 63.3 | 63.7 | 63.3 | 63.6 | 63.8 | 63.2 | 64.0 | 63.0 | 63.1 | 64.2 | 63.7 | 63.8 | 63.1 | 63.8 | 63.5 | 62.5 | | 65.0 |
| MN567045.1 Arabuta Snake virus-1 | 41.4 | 54.4 | 63.8 | 63.2 | 63.0 | 63.2 | 62.9 | 61.8 | 62.8 | 62.6 | 62.9 | 62.3 | 62.5 | 62.3 | 63.8 | 63.1 | 64.2 | 62.6 | 63.9 | 62.9 | 63.4 | 63.6 | 62.9 | 63.5 | |

**Supplementary Figure 5. Pairwise identity heatmap of reptarenavirus L sequences.** Pairwise identity heatmap of reptarenavirus L sequences are shown. Upper and lower triangles indicate nucleotide and amino acid identities, respectively.
