## Supplementary Information for "Identification of a novel and divergent reptarenavirus in an Amazon coral snake (*Micrurus spixii* [Wagler, 1824])"

**Full Methods**

***Detection of arenavirus-like contigs***

RNA-seq data were downloaded from NCBI SRA [1] and preprocessed using fastp 0.23.2 [2] with the default settings. The preprocessed reads were mapped to the genome of *Micrurus fulvius* (Eastern coral snake, ASM4626986v1) using HISAT2 2.2.1 [3] with the default settings, and unmapped reads were extracted by SAMtools [4] to remove host-derived reads. Because reference genomes for the species in which ACSV-1-derived reads were detected was not available, we used the genome of *M. fulvius*, which is closely related to that species according to TimeTree 5 [5]. The extracted unmapped reads were then assembled using SPAdes v3.15.5 [6], and contigs longer than 250 nucleotides were retained using SeqKit v2.3.0 [7].

The obtained contigs were subjected to sequence similarity searches against *Arenaviridae* [taxid:11617] using BLASTx 2.15.0+ [8] with the options "-word_size 2, -evalue 1e-4, and -max_target_seqs 5". Open reading frames (ORFs) in the contigs were translated and analyzed using the BLASTp web server (v2.17.0+; database: ClusteredNR; other parameters were set to default). Based on these results, contigs apparently derived from endogenous retroviruses or host transcripts were excluded, and only contigs whose best hit was to a reptarenavirus were used for subsequent analyses.

To evaluate the quality of the obtained reptarenavirus-like contigs, the original RNA-seq reads were mapped back to the corresponding contigs using HISAT2 v2.2.1 with the "-k 1" option. Read depth at each position was calculated using SAMtools. Positions supported by five or more reads were regarded as reliable in this study. Low-confidence regions were trimmed from the contigs. Furthermore, because the RNA-seq datasets (DRR089660–DRR089665) were generated using RACE-seq [9], adapter-derived sequences were trimmed from the contig ends.

***Annotation of ACSV-1 contigs***

ORFs longer than 1,000 nucleotides in the virus-like contigs were identified using Geneious Prime (Biomatters; https://www.geneious.com). In addition, for contigs containing L gene-like sequences,

ORFs longer than 100 nucleotides were also searched to identify the Z gene. For contigs containing the putative Z gene, ORFs longer than 200 nucleotides were identified. The predicted amino acid sequences were used as queries for BLASTp searches using the BLASTp web server (v2.17.0+; database: nr; organism: RNA viruses and retroviruses [taxid:2559587]; other parameters were set to default).

For the putative G protein, signal peptide, cleavage site, and transmembrane region were predicted by SignalP 6.0 [10] and Deep TMHMM 1.0 [11].

***Comprehensive detection of ACSV-1 in public RNA-seq datasets***

To detect ACSV-1 infection, 2,041 publicly available RNA-seq datasets from the suborder Serpentes (taxid: 8570) (Table S5) were mapped to the ACSV-1 contigs using Magic-BLAST v1.7.2 [1] with default settings. The numbers of mapped reads and per-base read depths were calculated using SAMtools. Read-depth profiles were then inspected manually, and datasets showing aberrant mapping patterns (e.g., a large number of reads mapping exclusively to a specific region within the untranslated region) were classified as negative.

The above mapping analysis was also applied to Squamata (taxid: 8509) (Table S6).

***Phylogenetic analysis***

To infer the phylogenetic relationship within the family *Arenaviridae*, phylogenetic analysis based on L protein sequences was performed as follows. First, the putative amino acid sequence of the ACSV-1 L protein, together with those of *Reptarenavirus*, was used as queries in BLASTp searches using BLAST 2.17.0+ web server (Database: nr; Organism: *Orthornavirae* [taxid: 2732396]; Max target sequences: 1000; E-value threshold: 1e−4; Word size: 2). Additionally, representative sequences from the genera other than *Reptarenavirus* within the family *Arenaviridae* [12] were included. The retrieved sequences were clustered using CD-HIT [13] with the "-c 0.9" option, and sequences shorter than 1,800 amino acids were removed. When viruses with specific names were clustered with the virus named “unidentified Reptareanavirus,” the named virus was selected. The sequences were then aligned using MAFFT with the E-INS-i algorithm and default settings, and unambiguously aligned regions were trimmed with trimAl [14] with the “-strict” option. Based on the alignment, the phylogenetic tree was reconstructed by the maximum likelihood method with RAxML Next Generation 1.1.0 [15] using the LG+I+G4+F model chosen by ModelTest-NG [16].

To further infer the evolutionary relationships within the genus *Reptarenavirus*, the NP and GP genes were also used for phylogenetic analyses. BLASTp searches were conducted using the putative amino acid sequences of the ACSV-1 NP and GP proteins as queries (Database: nr, Organism: *Orthornavirae* [taxid:2732396], Max target sequences: 1000, E-value threshold: $10^{-4}$, Word size: 2) on the BLAST web server. The BLAST hits belonging to the genus *Reptarenavirus* were extracted. The ACSV-1 and the extracted reptarenavirus sequences were subjected to clustering using CD-HIT [7] with the “-c 0.95” option. When viruses with specific names were clustered with the virus named “unidentified reptareanavirus,” the named virus was selected. The clustered sequences were then aligned by MAFFT using the E-INS-i algorithm with default settings, and unambiguously aligned were trimmed by trimAl [14] with the “-strict” option. Based on the alignments, phylogenetic trees were reconstructed as described above. The LG+G4 and FLU+G4 models chosen by ModelTest-NG were used for the NP and GP trees, respectively.
